# Impact of *Bactrocera oleae* on the fungal microbiota of ripe olive drupes

**DOI:** 10.1101/342543

**Authors:** Ahmed Abdelfattah, David Ruano-Rosa, Santa O. Cacciola, Maria G. Li Destri Nicosia, Leonardo Schena

## Abstract

The olive fruit fly (OFF), *Bactrocera oleae* is the most devastating pest affecting olive fruit worldwide. Previous investigations have addressed the fungal microbiome associated with olive drupes or *B. oleae*, but the impact of the insect on fungal communities of olive fruit remains undescribed. In the present work, the fungal microbiome of olive drupes, infested and non-infested by the OFF, was investigated in four different localities and cultivars. Olive fruit fly infestations caused a general reduction of the fungal diversity, a higher quantity of the total DNA and an increase in taxa that remained unidentified or had unknown roles. The infestations led to imbalanced fungal communities with the growth of taxa that are usually outcompeted. While it was difficult to establish a cause-effect link between fly infestation and specific fungi, it is clear that the fly alters the natural microbial balance, especially the low abundant taxa. On the other hand, the most abundant ones, were not significantly influenced by the insect. In fact, despite the slight variation between the sampling locations, *Aureobasidium*, *Cladosporium*, and *Alternaria*, were the dominant genera, suggesting the existence of a typical olive fungal microbiome.

## Introduction

Olive (*Olea europaea* L.) is one of the most important cultivated crops on a global scale (1). The countries of the coastal areas of the Mediterranean Basin represent the typical olive belt with more than 10 million ha, accounting for about 80% of the world’s total olive cultivation area (2). This crop is threatened by several abiotic and biotic stresses mainly caused by insects, pathogens, and nematodes (3-5). Among those, the olive fruit fly (OFF) *Bactrocera oleae* (Rossi) is considered the most destructive key pest for olive production, and can be virtually found wherever *Olea* species are present (6). *Bactrocera oleae*, in contrast to other *Tephritidae Diptera*, is strictly monophagous, feeding exclusively on olive fruits, and has a unique ability to feed on unripe green fruits which contain high levels of phenolic compounds such as oleuropein. This ability is attributed to the high number of detoxifying genes activated during larvae feeding (7) as well as the presence of the symbionts such as ‘*Candidatus* Erwinia dacicola’ in their gut (8). The microbiome of *B. oleae* has been found to be essential for its survival in natural conditions and is likely to play an important role on the longevity, competitiveness, and ability of the insect to cope with biotic stresses (9). Although the source of *B. oleae* microbiome is unknown, some evidence suggests a vertical transmission from one generation to the next. Adult females of *B. oleae*, use their ovipositors to lay eggs inside olive drupes, and the hatched larvae form tunnels inside the fruit while feeding on the mesocarp. The physical damage, represented in fruit punctures and larvae feedings affect both crop yield and fruit and oil quality dramatically. In particular, *B. oleae* infestations affect phenolic content, fatty acids profile, and peroxide values of olive oil (10). As a response to *B. oleae* infestation, the plant produces different defensive compounds including phytohormones, volatile organic compounds, and defense proteins, among others (11). It has been also reported that the insect can facilitate secondary microbial infections caused by different microorganisms (10). Despite the fact that these secondary colorizers are likely to affect olive oil quality, little is currently known about the impact of olive fly infestations on olive microbial communities. The role of fungi is particularly relevant in this context due to their importance as plant pathogens(12), and capability to produce mycotoxins. Indeed, recent microbiome studies have mainly focused on bacterial and, to a lesser extent, fungal communities associated with *B. oleae* (13). Other investigations have focused on the fungal communities associated with different organs of the olive canopy including ripe drupes (14) but the impact of *B. oleae* on fungal communities has never been investigated. In the present study, the relationship between insect infestation and fungi has been analyzed by determining fungal diversity and total fungal DNA in infested and non-infested drupes from different cultivation regions in Italy.

## Materials and methods

### Sampling and DNA extraction from olive drupes

Samples, were collected in the middle of November 2015 from four commercial olive orchards of the cultivars Lection, Ottobratica, Carolea, and Ciciarello located in Monopoli (Puglia, Southern Italy), and in Mileto, Filadelfia and Maierato (Calabria, Southern Italy), respectively (Table 1).

**Table 1.** Source of infested and non-infested olive samples analyzed in the present study

| Locality | Cultivar | Age (years) | GPS coordinates |
| --- | --- | --- | --- |
| Monopoli, Bari | Leccino | 20 | 40°53'09.5"N 17°18'22.6"E |
| Mileto, Vibo Valentia | Ottobratica | 30 | 38°36'05.4"N 16°04'06.4"E |
| Filadelfia | Carolea | 30 | 38°47'33.9"N 16°16'55.1"E |
| Maierato | Ciciarello | 30 | 38°43'17.5"N 16°12'42.6"E |

From each location, 5 biological replicates of both infested and non-infested olives were randomly collected from an area of approximately 1 ha. Each biological replicate consisted of ten drupes. Collected olives were quickly returned to the laboratory in plastic containers and stored at 4 °C for no longer than 24 h prior to lyophilization (Labconco^®^ FreeZone 2.5). Before lyophilization the fruits’ mesocarp and exocarp were separated from the stone, which was discarded. Lyophilized samples were grounded in liquid nitrogen with sterile mortars and pestles, and total DNA was extracted from 80 mg of homogenate tissues as described by Mosca and co-workers (15). Concentration and quality of extracted DNA were assessed using Nanodrop spectrophotometer (Thermo Fisher Scientific Inc., USA). All samples were diluted to have a DNA concentration of 20 μg·μl^−1^.

### Quantification of total fungal DNA

Total fungal DNA was estimated by quantitative real-time PCR using the universal fungal primers ITS3_KYO2-ITS4 targeting the fungal ITS2 region of the rDNA (16) and the SYBR chemistry (Applied Biosystems, USA). To construct a representative standard curve, total DNA was extracted from a pure culture of *Colletotrichum acutatum sensu stricto* as described by Schena & Cooke (17), quantified using a Nanodrop spectrophotometer, and serially diluted with sterile water to yield final DNA concentrations ranging from 10 ng/μl to 100 fg/μl. PCR reactions were performed in a total volume of 20 µl containing 10 μl SYBR Green PCR Master Mix, 300 nM of each primer, and 2 μl of DNA template (20 ng/μl). All amplifications were performed in triplicate and water replaced template DNA in negative control reactions. Amplifications were performed and monitored using the StepOnePlus Real-Time PCR System (Applied Biosystems, USA) and data acquisition, and analysis was made using the supplied software according to the manufacturer’s instructions. A standard curve was generated by plotting the DNA quantities against the corresponding Cq value. Correlation coefficients, mathematical functions, and PCR efficiencies were calculated and visualized using StepOne software.

The total quantity of fungal DNA in olive samples was quantified by amplifying 2 µl of DNA extracted from each biological replicate. The Cq value obtained for each biological replicate was an average of the three technical replicates. To take into account small variations in sample size and efficiency of extraction and amplification, the data were normalized using a qPCR method based on two universal primers and a probe (17). Specifically, for each biological replicate, Cq values obtained with fungal universal primers were corrected against Cq values of the universal method (Cq-u) by multiplying the uncorrected Cq values by the ratio of the average of Cq-u and the Cq-u of each specific sub-sample. The concentration of total fungal DNA in unknown samples was extrapolated using corrected Cq values and the specific mathematical functions (regression solutions) determined for the standard curves. The total DNA concentration was expressed as ng of fungal DNA per g of fresh olive tissue.

Data were submitted to statistical analysis using the total DNA quantity as dependent variable and the location of sampling or the OFF infestations as fixed factors using SPSS software (IBM Corp. Released 2013. IBM SPSS Statistics for Windows, Version 22.0. Armonk, NY: IBM Corp.). In the first case, analyses were separately performed for infested and non-infested olives and means were compared by a two-sided test pairwise comparison. In the latter case, analyses were performed for each locality by the ANOVA. Furthermore, a two-factor essay (general linear model) was performed in order to determine the effect of location and fly infestations and their interactions on the total fungal quantity.

## Metabarcoding analyses

### Library preparation and sequencing

DNA extracts were amplified using the universal fungal primers ITS3_KYO2-ITS4, modified by adding the adapters needed for Illumina MiSeq sequencing. PCR reactions were conducted in a total volume of 25 μl, consisting of 1 μl of DNA (50 μg) or nuclease-free water (QIAGEN, Valencia, CA, USA) as negative control, 12.5 μl of KAPA HiFi HotStart ReadyMix (Kapa Biosystems, Wilmington, MA, USA) and 1.5 μl of each primer (10 μM). Amplification was performed in an Eppendorf Vapo Protect Mastercycler pro S (Eppendorf, Germany) as follows: 3 min at 98 °C followed by 30 cycles of 30 s at 95 °C, 30 s at 50 °C and 30 s at 72 °C. PCR products were purified using the Agencourt Ampure XP kit (BECKMAN COULTER-Beverly, Massachusetts), multiplexed using the Nextera XT indexing kit (Illumina, Inc., San Diego, CA), and purified again with the Agencourt Ampure XP kit. All purification and amplification reactions were performed according to the manufacturer’s instructions. The concentration of purified amplicons was measured using Qubit 3.0 Fluorometer (Thermo Fisher Scientific, Waltham, Massachusetts) and normalized to a common concentration of 20 ng/μl. Normalized amplicons were pooled by adding 10 μl of each product and kept at −20 °C until sequencing. Sequencing was performed on an Illumina MiSeq platform by loading 4 nM of denatured DNA into V3 MiSeq flow cell for 600 cycles (300 × 2) according to Illumina standard protocol.

### Analysis of metabarcoding data

Raw Illumina data were processed using the Trimmomatic V0.32 (18) and PANDAseq assembler V2.11 (19) to eliminate low quality reads and merge paired-end sequences with a minimum of 100 bp read length. The downstream analyses were done using QIIME 1.9.1 pipeline (20) as described by Abdelfattah and co-workers (21). The operational taxonomic Units (OTUs) table was normalized by rarefaction to an even sequencing depth in order to remove sample heterogeneity.

The rarefied OTU table was used to calculate alpha diversity indices including Observed Species (*Sobs*) and Shannon metrics and to plot taxa relative abundances (RA). MetagenomeSeq’s cumulative sum scaling (CSS) (22) was used as a normalization method for other downstream analyses. Alpha diversities were compared based on a two-sample t-test using non-parametric (Monte Carlo) methods and 999 Monte Carlo permutations (999). The CSS normalized OTU table was used to calculate the β-diversity using the Bray Curtis metrics (23) and construct the principal coordinate analysis (PCoA) plots (24). To evaluate the significance of differences in RAs of detected taxa, the most abundant ones (≥ 0.1%) were compared using the Kruskal-Wallis test (25). In all tests, significance was determined using 1000 Monte Carlo permutations, and the BenjaminiHochberg (FDR) corrections were used to adjust the calculated p-values.

### Data Availability

The datasets generated during the current study were deposited and are available at the National Center for Biotechnology Information (NCBI), Sequence Read Archive (SRA), under the accession number PRJNA427133 (www.ncbi.nlm.nih.gov/bioproject/427133). Other data generated or analyzed during this study are included in this published article.

## Results

### Construction of a standard curve for quantitative analyses

Total fungal DNA was estimated by quantitative real-time PCR using the universal fungal primers. Standard curve generated using serial dilutions of DNA of *C. acutatum* revealed a linear correlation over six orders of magnitude (from 10 ng to 100 fg) with a correlation coefficient (R2) of 0.999. A specific linear equation (y=20.31-3.30x) was established and utilized to quantify the total fungal DNA in olive drupes.

### Fungal diversity and richness

After quality filtering, Illumina high throughput sequencing resulted in a total of 3,019,642 reads, with a mean of 86,275 reads per sample. Overall, these sequences were assigned to 1,428 OTUs (97% similarity threshold) with a mean of 123 OTUs per biological replicate. According to the taxonomy assignments, the OTUs were assigned to the phyla *Ascomycota* (89.9%), *Basidiomycota* (9.7%), and a very small fraction to *Zygomycota* (0.01%) and unidentified fungi (0.3%) (Fig. 1).

**Figure 1.**
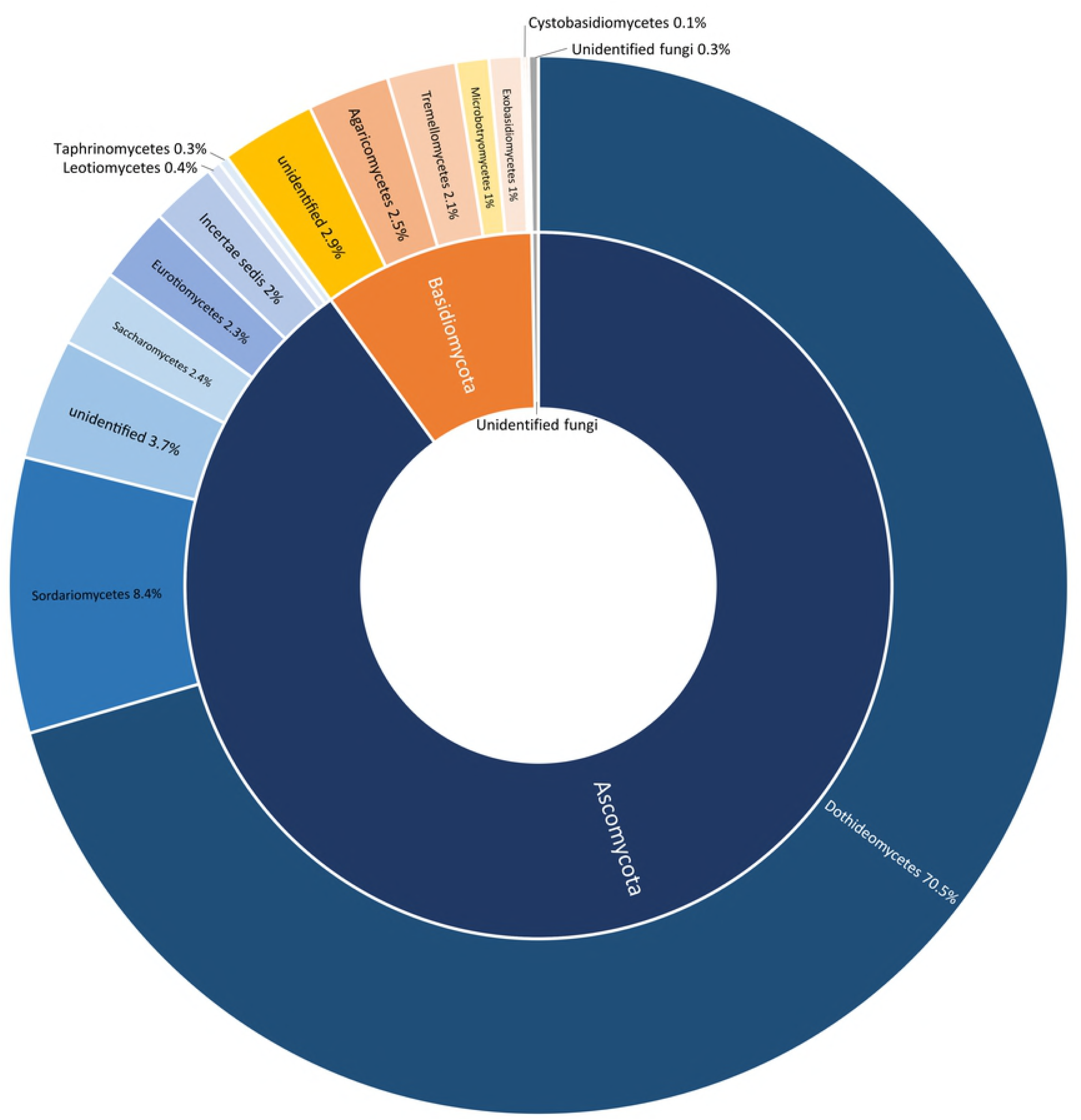
Sunburst chart showing the total relative abundance of fungal phyla (interior circle) and classes (exterior circle) overall detected in infested and non-infested olives in the four investigated localities.

Most *Ascomycota* were represented in the classes *Dothideomycetes* (70.5%) and *Sordariomycetes* (8.4%), whereas the majority of the *Basidiomycota* was represented by an unidentified class (2.9%), *Agaricomycetes* (2.5%), and *Tremellomycetes* (2.1%) (Fig. 1). At the genus level, the most abundant genera were *Aureobasidium* (30.6%), *Cladosporium* (17.2%), *Alternaria* (11.8%), *Colletotrichum* (4.5%), Unidentified Ascomycota (3.7%), and *Pseudocercospora* (3.6%) (Fig. 2; Fig. 6).

**Figure 2.**
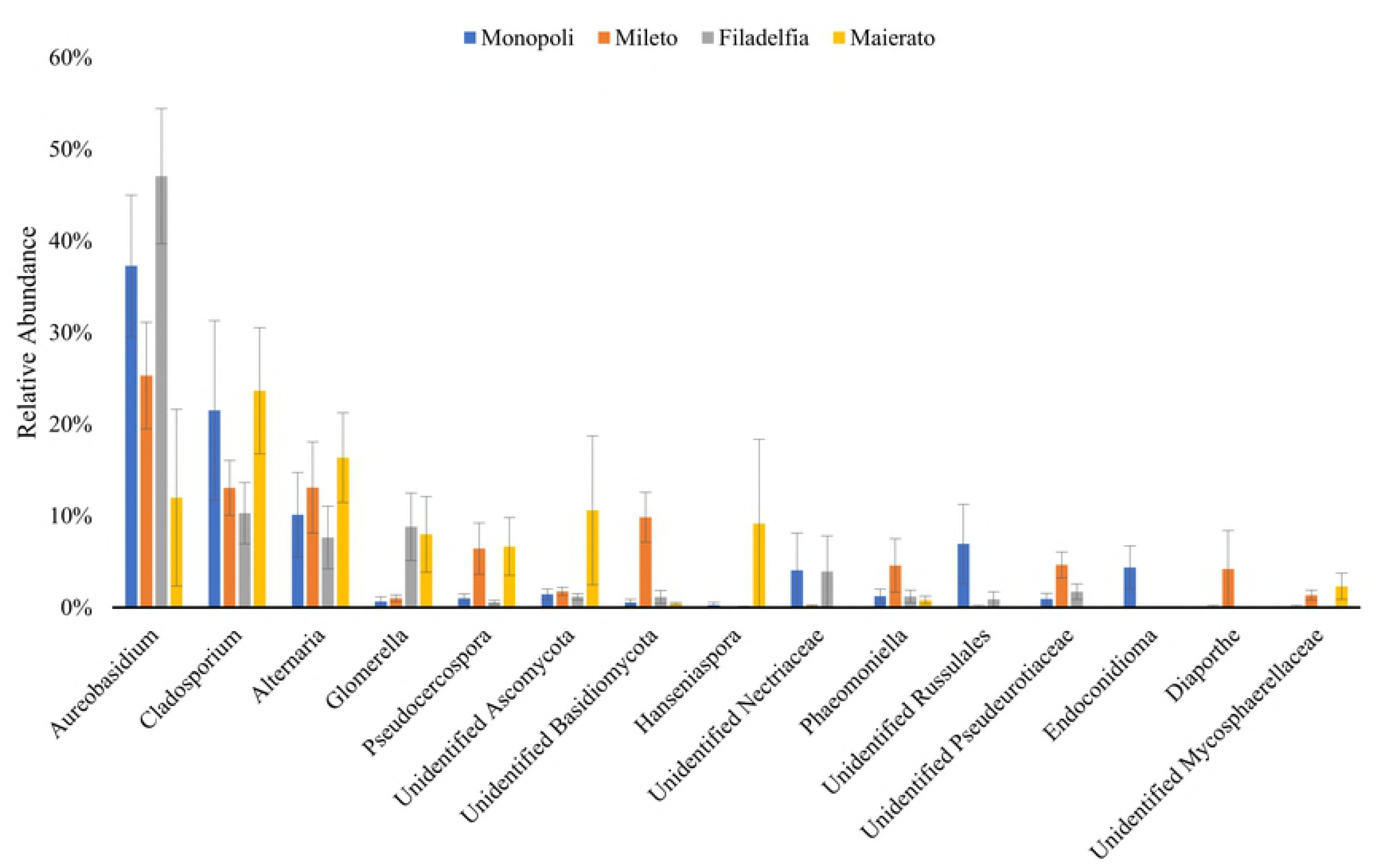
Relative abundances of the dominant fungal genera (RA ≥1%) detected in infested and non-infested olives in Monopoli, Mileto, Filadelfia, and Maierato. The error bars were calculated by dividing the standard deviation on the square root of the sample size.

### Comparison of locations

Quantitative real-time PCR analyses of olive drupes from four commercial orchards located in Monopoli, Mileto, Filadelfia and Maierato, revealed some significant differences in the total quantity of fungal DNA according to the investigated locations (Table 1; Fig. 3).

**Figure 3.**
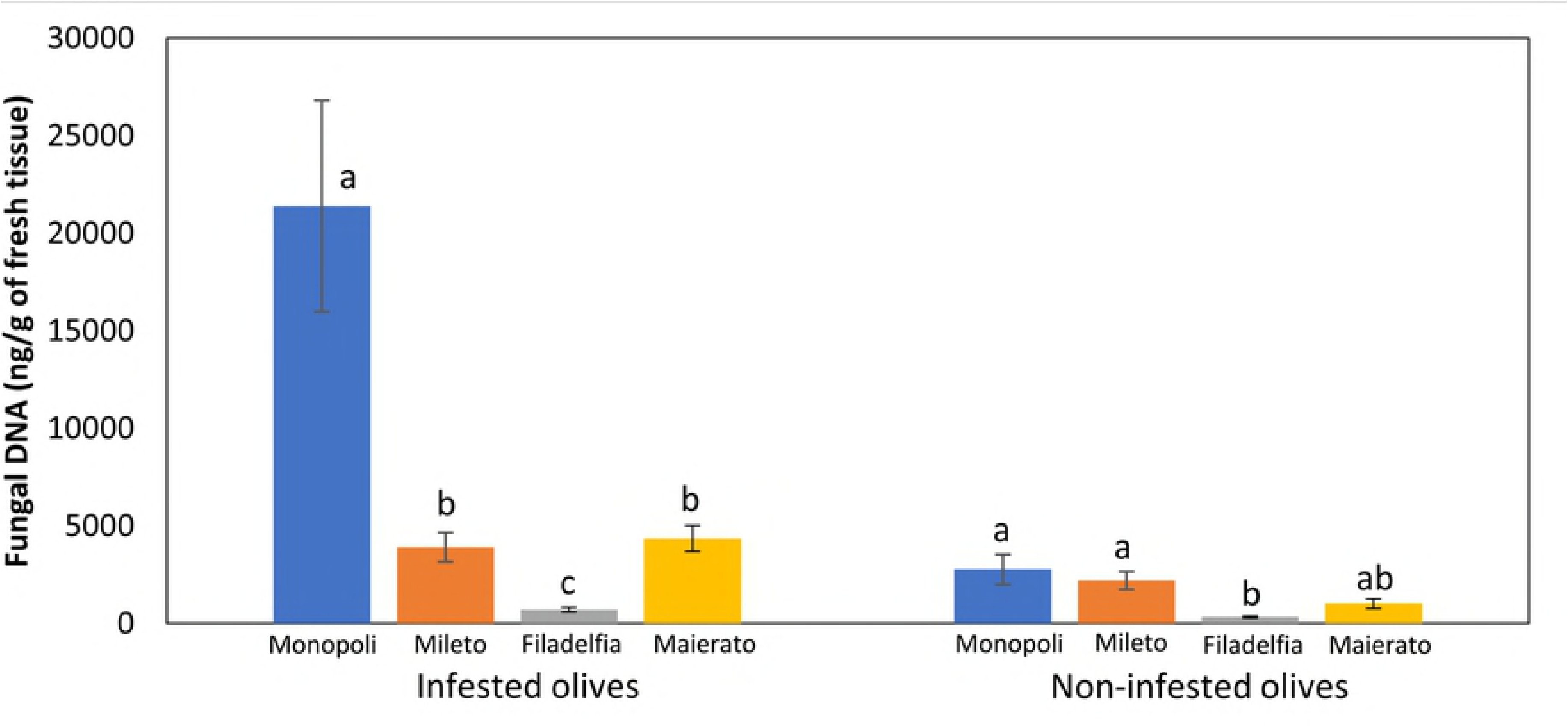
Estimated concentration of fungal DNA in olive samples collected in Monopoli, Mileto, Filadelfia and Maierato. Independent statistical analyses were performed for infested (left side) and non-infested (right side) olive samples. Different letters indicate significantly different values according to 2-sided tests (P≤ 0.05). Bars indicate standard errors of the means.

However, differences were more remarkable in infested olives as compared to the non-infested ones. In the first case, significant differences were revealed between most samples since the total quantity of detected DNA per g of fresh tissue ranged between 21400 (Monopoli) and 717 ng (Filadelfia). In the latter case, most samples were not significantly different and total DNA ranged from 2787 ng and 341 ng (Fig. 3). According to metabarcoding analyses, a consistent fungal community was associated with olive drupes in all investigated localities since three genera (*Aureobasidium*, *Cladosporium* and *Alternaria*) cumulatively accounting for approximately 60% of the detected sequences, were always the prevalent fungi (Fig. 2).

On the other hand, the number of observed species (*sobs* index) differed significantly in the four sampling locations, with the exception of Maierato and Filadelfia (Table 2). A lower level of discrimination was provided by the Shannon index, since significant differences were only revealed between Mileto and both Filadelfia and Monopoli (Table 2).

**Table 2.** Comparison of the alpha diversity of the fungal communities associated to olive drupes in the four investigated localities (Monopoli, Mileto, Filadelfia, and Maierato) according to Shannon and observed species (*sobs*) indexes. P values of the pair-wise comparisons were determined using student-t-test and 999 Monte Carlo permutations. Both Shannon and Sobs Mean values are reported in bold for each investigated locality.

|  | *Shannon* |  |  |  | *sobs* |  |  |  |
| --- | --- | --- | --- | --- | --- | --- | --- | --- |
|  | Monopoli | Mileto | Filadelfia | Maierato | Monopoli | Mileto | Filadelfia | Maierato |
| Monopoli | **2.08** | 0.04 | 0.47 | 0.68 | **28.56** | 0.015 | 0.024 | 0.006 |
| Mileto |  | **3.23** | 0.03 | 0.05 |  | **63.90** | 0.024 | 0.045 |
| Filadelfia |  |  | **2.34** | 0.81 |  |  | **36.91** | 0.115 |
| Maierato |  |  |  | **2.27** |  |  |  | **42.74** |

In particular, significant differences among investigated localities were revealed for 21 taxonomic entities that comprised both abundantly detected taxa such as *Aureobasidium*, *Colletotrichum* and *Pseudocercospora* and much less abundant taxa (Table 3).

**Table 3.** List of fungal taxa characterized by a significantly different relative abundance (RA) between the four investigated locations. P values were calculated using nonparametric test of Kruskal-Wallis through 1000 permutations. FDR P are the P value corrected by the Benjamini-Hochberg FDR procedure.

| Detected taxa | FDR P | Relative abundance (%) |  |  |  |
| --- | --- | --- | --- | --- | --- |
|  |  | Monopoli | Mileto | Filadelfia | Maierato |
| *Aureobasidium* | 0.01232 | 39.11 | 26.66 | 48.91 | 12.18 |
| *Colletotrichum* | 0.00887 | 0.18 | 1.05 | 9.14 | 8.36 |
| *Pseudocercospora* | 0.04381 | 0.69 | 6.88 | 0.61 | 6.91 |
| Unidentified Basidiomycota | 0.00621 | 0.27 | 10.80 | 1.18 | 0.45 |
| Unidentified Pseudeurotiaceae | 0.00156 | 0.86 | 5.12 | 1.80 | 0.00 |
| *Endoconidioma* | 0.00029 | 4.93 | 0.04 | 0.03 | 0.00 |
| *Diaporthe* | 0.00533 | 0.00 | 4.24 | 0.03 | 0.00 |
| Unidentified Mycosphaerellaceae | 0.00156 | 0.00 | 1.42 | 0.00 | 2.44 |
| *Quambalaria* | 0.00621 | 0.10 | 0.00 | 2.96 | 0.00 |
| *Fusicladium* | 0.00029 | 0.00 | 0.01 | 0.91 | 2.05 |
| *Sporobolomyces* | 0.00666 | 1.38 | 0.56 | 0.80 | 0.00 |
| *Botryosphaeria* | 0.00404 | 0.00 | 2.13 | 0.00 | 0.00 |
| *Dioszegia* | 0.00263 | 0.00 | 0.01 | 1.87 | 0.00 |
| *Stemphylium* | 0.00130 | 0.09 | 0.21 | 0.00 | 1.44 |
| *Plenodomus* | 0.01232 | 0.00 | 0.00 | 1.52 | 0.00 |
| Unidentified Fungi | 0.02754 | 1.01 | 0.03 | 0.25 | 0.00 |
| *Taphrina* | 0.00072 | 0.00 | 1.01 | 0.00 | 0.06 |
| Unidentified Tremellales | 0.00079 | 0.00 | 0.75 | 0.00 | 0.00 |
| Unidentified Tremellomycetes | 0.00621 | 0.56 | 0.00 | 0.00 | 0.00 |
| *Tilletiopsis* | 0.03415 | 0.00 | 0.01 | 0.00 | 0.46 |
| *Toxicocladosporium* | 0.00031 | 0.00 | 0.00 | 0.00 | 0.43 |

Furthermore, the analysis of beta diversity showed that the fungal community significantly differed between all the sampling locations (Table 4).

**Table 4.** Comparison of alpha and beta diversity in infested and non-infested olive drupes collected in Monopoli, Mileto, Filadelfia, or Maierato, according to Shannon and observed species (*sobs*) indexes. P-values were determined using 999 Monte Carlo permutations.

|  | Observed species (*sobs*) |  | Shannon |  | P value |  |  |
| --- | --- | --- | --- | --- | --- | --- | --- |
|  | Infested | Non-infested | Infested | Non-infested | Sobs | Shannon | Beta diversity |
| Monopoli | 25.8 | 31.3 | 2.11 | 2.05 | 0.18 | 0.95 | 0.033 |
| Mileto | 38.2 | 84.5 | 2.50 | 3.82 | 0.07 | 0.06 | 0.005 |
| Filadelfia | 39.7 | 33.4 | 2.30 | 2.38 | 0.25 | 0.90 | 0.016 |
| Maierato | 39.0 | 47.4 | 1.91 | 2.73 | 0.19 | 0.27 | 0.029 |
| All locations | 36.1 | 51.2 | 2.19 | 2.80 | 0.025 | 0.02 | 0.001 |

These differences were particularly evident when beta diversity results were visualized on a PCoA plot since samples from all localities segregated separately (Fig. 4).

**Figure 4.**
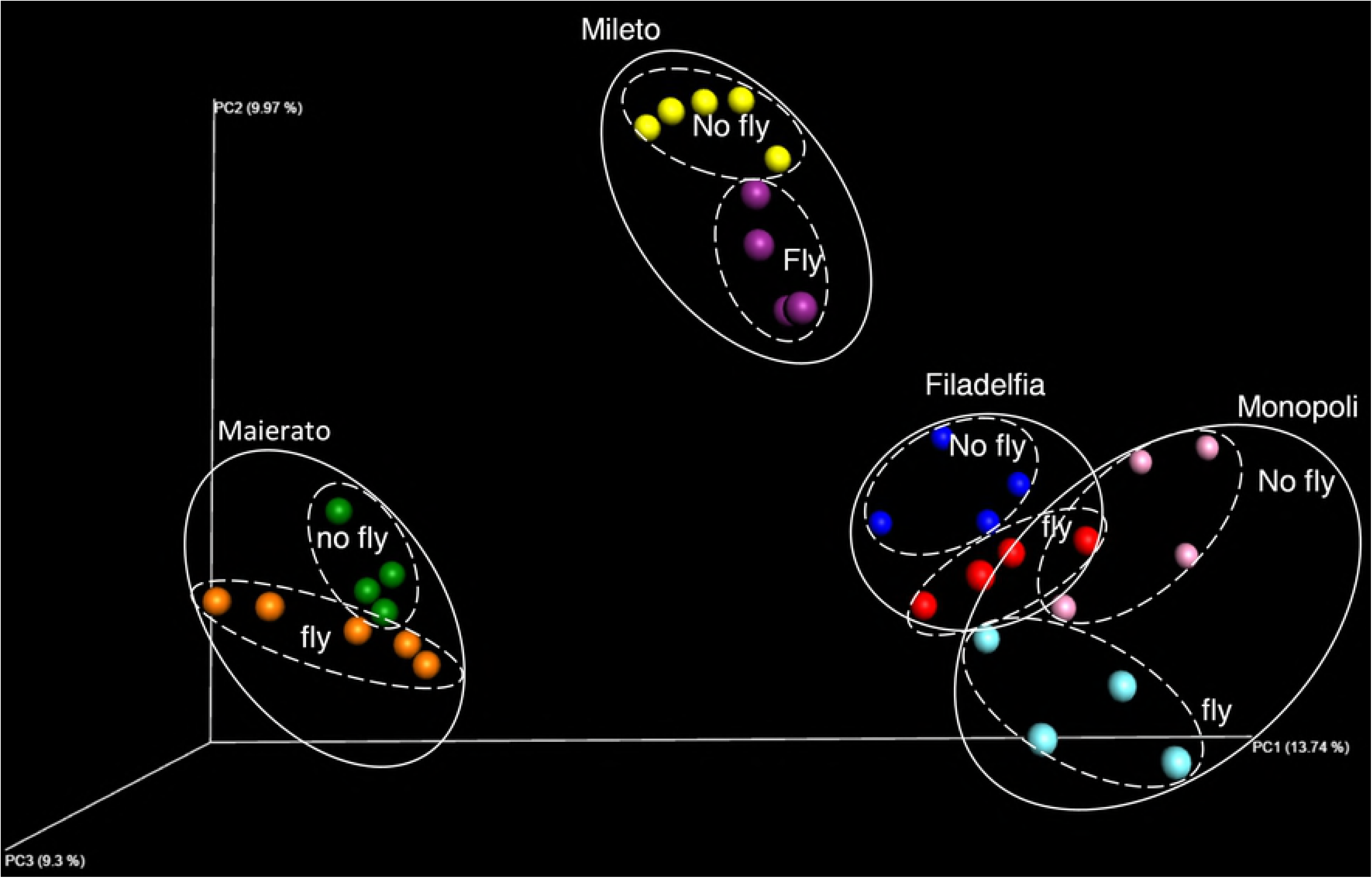
Comparison of total fungal DNA in infested and non-infested olive drupes in the four investigated localities (Monopoli, Mileto, Filadelfia and Maierato). Independent statistical analyses were performed for each locality. Different letters indicate significantly different values according to ANOVA univariate analysis of variance (P ≤ 0.05). Bars indicate standard errors of the means.

### Comparison of infested and non-infested olives

The total quantity of fungal DNA was higher in infested olives as compared to the non-infested ones in all localities. In particular, total fungal DNA increased by 87.0, 43.6, 52.4, and 76.5%, in Monopoli, Mileto, Filadelfia, and Maierato, respectively, although differences were statistically significant only in Monopoli and Maierato (Fig. 5).

**Figure 5.**
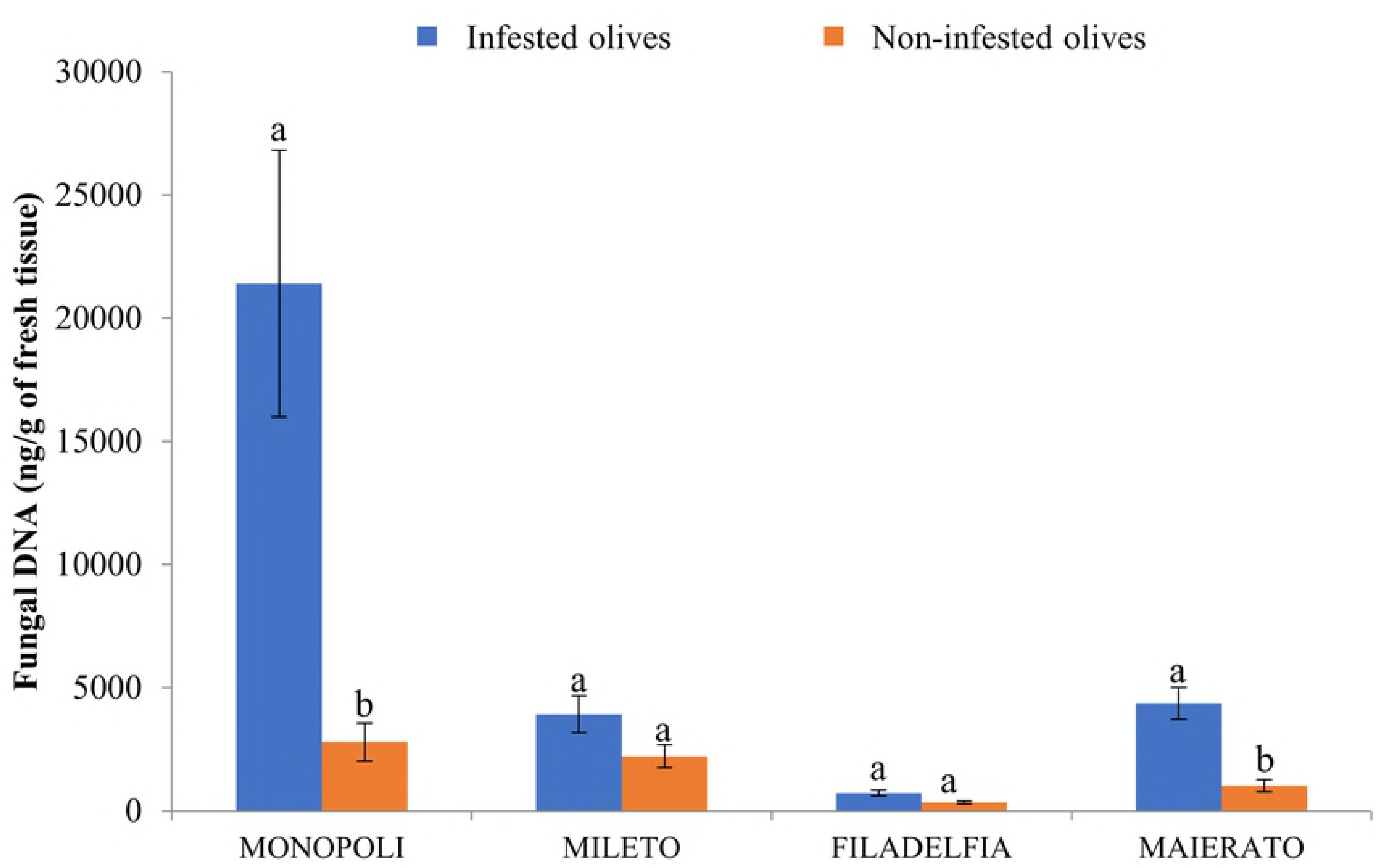
Principal coordinate analysis (PCoA) of the fungal communities associated with olive drupes infested (fly) or non-infested (no fly) by the olive fruit fly Bactrocera oleae and collected in the four studied localities (Monopli, Mileto, Filadelfia or Maierato).

The ANOVA two-factor essay confirmed that the fungal DNA quantity varied depending on the fly puncture (F= 18,541; df=1; P<0.001), the site (F= 13,474; df=3; P<0.001), and the interaction of both factors (F=9,254; df=3; P<0.001).

A significant impact of the OFF on the composition of the fungal community was confirmed by metagenomic analyses. Regardless of the location, the number of observed species was significantly higher in non-infested fruits with a mean of 49.1 OTUs compared to 35.7 OTUs in infested fruits (P =0.025). Similarly, healthy fruits had a higher Shannon index value (2.75) compared to infested fruits (2.2) (P =0.020). Despite these differences, both the number of observed species and Shannon index were not statistically significant when infested and non-infested fruits were compared individually (Table 4). Interestingly, most of the taxa with significant different RA in infested and non-infested olives had a quite low RA and/or remained unidentified (Table 5). On the other hand, the most abundant taxa such as *Aureobasidium*, *Cladosporium* and *Alternaria*, did not differ significantly in their RAs between infested and non-infested olives (Table 5).

**Table 5.** List of taza with a significant different relative abundance (RA) in infected and non-infected olive drupes, in the four investigated locations.

| Location | Taxa | P-value | Relative abundance (%) |  |
| --- | --- | --- | --- | --- |
|  |  |  | Non-infested | Infested |
| Monopoli | Unidentified *Tremellomycetes* | 0.0139 | 1.12 | 0.00 |
|  | Unidentified *Pseudeurotiaceae* | 0.0180 | 1.72 | 0.01 |
|  | Unidentified *Russulales* | 0.0180 | 0.01 | 15.20 |
|  | *Quambalaria* | 0.0180 | 0.01 | 0.20 |
|  | Unidentified *Basidiomycota* | 0.0180 | 0.53 | 0.01 |
|  | Unidentified Fungi | 0.0180 | 2.01 | 0.01 |
|  | Unidentified *Sporobolomyces* | 0.0202 | 2.74 | 0.02 |
| Mileto | *Sporobolomyces* | 0.0105 | 1.01 | 0.00 |
|  | *Phaeoramularia* | 0.0127 | 2.46 | 0.08 |
|  | *Colletotrichum* | 0.0127 | 1.88 | 0.01 |
|  | Unidentified *Mycosphaerellaceae* | 0.0143 | 2.53 | 0.04 |
|  | Unidentified *Pseudeurotiaceae* | 0.0143 | 8.64 | 0.72 |
|  | *Taphrina* | 0.0143 | 1.76 | 0.06 |
|  | Unidentified *Basidiomycota* | 0.0143 | 17.81 | 2.04 |
|  | Unidentified *Nectriaceae* | 0.0318 | 0.32 | 0.00 |
|  | *Phaeomoniella* | 0.0491 | 0.65 | 10.11 |
| Filadelfia | Unidentified *Dothideomycetes* | 0.0127 | 1.56 | 0.03 |
|  | *Cryptococcus* | 0.0139 | 0.03 | 0.45 |
|  | Unidentified *Basidiomycota* | 0.0139 | 0.08 | 2.06 |
|  | *Plenodomus* | 0.0318 | 0.00 | 2.74 |
| Maierato | *Toxicocladosporium* | 0.0139 | 0.04 | 0.92 |
|  | *Fusicladium* | 0.0275 | 0.62 | 3.84 |

In particular, most abundant taxa showed different and inconsistent trends in infested and non-infested olives according to the sampling localization (Fig. 6). For instance, the genus *Aureobasidium* had a higher RA in infested olives in Filadelfia, Mileto and Maierato and lower abundance in Monopoli. The genus *Cladosporium* was more abundant in infested olives in Filadelfia and Mileto and less abundant in Monopoli and Maierato. The genus *Alternaria* was more abundant in infested olives in Monopoli and more abundant in non-infested drupes in the other sampling sites (Fig. 6).

**Figure 6.**
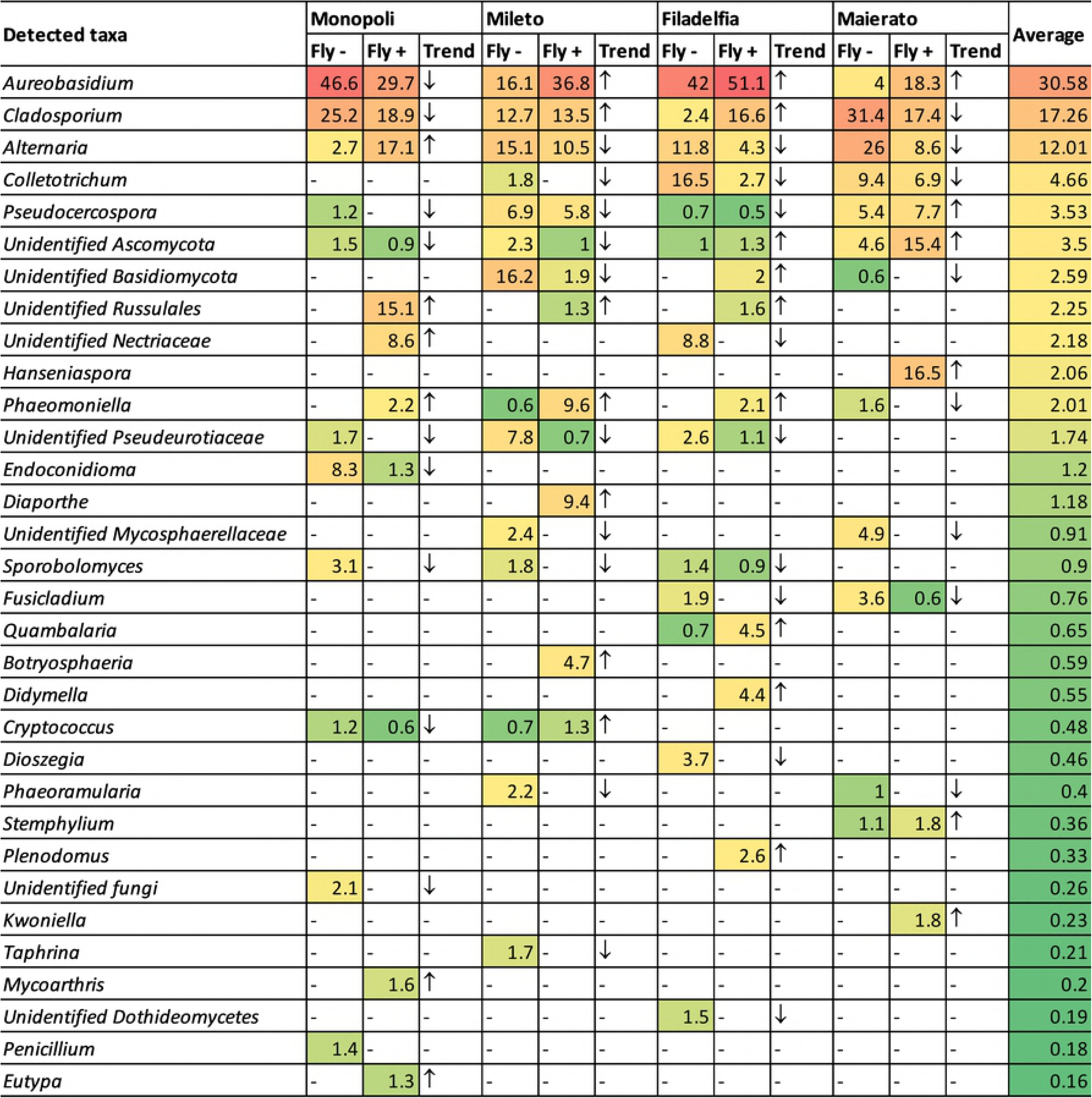
Heatmap highlighting the differences in the taxa detected in infested (Fly +) and non-infested (Fly -) olives from four different localities with an overall relative abundance of at least 0.16%. Arrows indicate the trend of each taxa in infested olives as compared to the non-infested ones.

A clearer picture of the impact of the OFF on olive fungal community was achieved through the beta diversity analysis. In fact, *B. oleae* infestations caused a significant change in the fungal community structure in all investigated locations (Filadelfia P = 0.016, Maierato P = 0.029, Mileto P = 0.005, and Monopoli P = 0.033) and differences were confirmed by cumulatively analyzing drupes from the four sampling sites (P=0.001) (Table 4). In agreement with the beta diversity analyses, the PCoA visualization of results showed a separate clustering of infested and non-infested samples in all the investigated localities (Fig. 4)

## Discussion

The present work enabled the detailed investigation of the impact of *B. oleae* on the structure of the fungal community of the olive carposphere. Interestingly, OFF infestations resulted in a general reduction in fungal diversity, but also in a significant increase in the total quantity of fungi. Although these results were shared by all investigated localities, the impact of OFF on the fungal community structure varied in the investigated localities. It appeared that flies had a unique influence in each locality in relation to different environmental conditions, cultivars, and management practices. Another important factor that was likely to influence results was the average lapse of time between the fly puncture and sampling. Indeed, olives collected in Monopoli and Maierato were characterized by older infestations and this may explain the differences in terms of total quantity of fungal DNA since fungi had more time for the colonization of injured tissues.

A significant impact of *B. oleae* infestation on the structure of fungal communities was confirmed by beta diversity analyses since they showed a clear separation of infested and non-infested samples in all investigated locations. Interestingly, differences were more noticeable in taxa with a quite low RA while abundant taxa such as *Aureobasidium*, *Cladosporium*, and *Alternaria*, did not differ significantly and showed inconsistent trends in the different sampling locations. Overall, the obtained data suggest that OFF infestations led to imbalanced fungal communities. Indeed, while it is difficult to make concrete speculations on the role of specific fungal genera in infested fruits, the reduced microbial diversity and the higher quantity of fungal DNA support the hypothesis of an altered equilibrium among taxa. The abundant presence of taxa with unknown roles as well as unidentified taxa indicate that the infestation of OFF may have facilitated the colonization of the fruit by alien species and/or favored the growth of taxa that are usually outcompeted. Similar phenomena, named dysbiosis, occur in humans when one or more factor is able to change the structure of the microbial community (26). Dysbiosis is usually used to describe a condition where the, normally, most abundant taxa are less represented and are being replaced by less common ones (27). There is no single cause of dysbiosis, even though it has been associated with diseases and inappropriate drugs or diet (28). It is not far-fetched to hypothesize that the infestation by OFF may have caused a similar condition in the investigated olives. Although dysbiosis has never been reported in plants, the consistency of results obtained in four different localities characterized by different environmental conditions, cultivars and management practices, supports the presence of such phenomena. While it is difficult to establish a cause-effect link between fly and specific fungal taxa, it is clear that the fly alters the natural microbial balance and this may be linked to the reduced oil quality made from infested fruits. Indeed, fungal infections are known to change the oil quality by increasing its acidity or peroxide values (29).

The consensus of the olive community composition described herein, including the proportion of the fungal taxa at all the taxonomical levels (from phyla to genera) is in strong agreement with the previous description of the olive fungal community; being dominated by fungal genera such as *Aureobasidium*, *Cladosporium*, and *Alternaria* (14). The stability of the olive fungal community in the investigated samples in the present and in a previous study (14), and how they differ remarkably from other hosts (21) (30) (21), confirms that the microbial community is determined primarily by its host identity. Interestingly, non-pathogenic fungal species such as *Aureobasidium pullulans* were largely prevalent in all localities. The abundant presence of this species is remarkable since this yeast-like fungus is well-known for its antimicrobial activity against a wide range of microbes (31-33) and has been used as biological control agent against several plant diseases (34-36). Regardless of the hypothetical role of *Aureobasidium*, little is known about the actual ecological reasons for its natural prevalence on olive plants as compared to other plant species.

Despite the characteristic fungal community associated with olives, in the present study it was difficult to determine a precise list of fungal taxa that defines infested and non-infested olives, also because some significant variations were observed between the investigated locations/cultivars. It is likely that these differences were the results of several factors including environmental conditions, cultivars, and management practices. In fact, previous studies have showed that the geographical location, among other factors such as the management practices, can influence the fungal community of the host (21, 37). Indeed, it has been proven that different organs, and even different parts of the same organ can significantly impact the microbial composition (21).

Interestingly, the majority of olive fungal community, described herein and in the previous study (14) overlap with those found on olive fruit fly (13). This is the first observation to document two different species, especially from two different kingdoms, to have a very similar fungal microbial community. In agreement with our results, the presence of several bacterial species common to both *B. oleae* and olive trees have already been reported (38). Due to the limited information currently available on this subject, it is difficult to make concrete speculations about the significance of our observation. However, since *B. oleae* is a host-specific insect, which spends most of its life cycle in association with its host tissue (10), the observed similarities may indicate a shared evolutionary process.

## Acknowledgement

We thank A. Davies for the revision of the English text and Dr. Anna Maria Cichello for her help during sampling. This research was funded by The Italian Ministry of Education, University and Research with grant “Modelli sostenibili e nuove tecnologie per la valorizzazione delle olive e dell’olio extravergine di oliva prodotto in Calabria” - PON Ricerca e competitività 2007-2013 (PON03PE_00090_02).

